# dadi.CUDA: Accelerating population genetic inference with Graphics Processing Units

**DOI:** 10.1101/2020.07.30.229336

**Authors:** Ryan N. Gutenkunst

## Abstract

Extracting insight from population genetic data often demands computationally intensive modeling. dadi is a popular program for fitting models of demographic history and natural selection to such data. Here, I show that running dadi on a Graphics Processing Unit (GPU) can speed computation by orders of magnitude compared to the CPU implementation, with minimal user burden. This speed increase enables the analysis of more complex models, which motivated the extension of dadi to four- and five-population models. Remarkably, dadi performs almost as well on inexpensive consumer-grade GPUs as on expensive server-grade GPUs. GPU computing thus offers large and accessible benefits to the community of dadi users. This functionality is available in dadi version 2.1.0, https://bitbucket.org/gutenkunstlab/dadi/.

---

Population genetic data contain much information about the history of the sampled populations, but computationally intensive modeling is often necessary to extract that information. dadi is widely used for inferring models of demographic history (Gutenkunst et al., 2009) and natural selection (Kim et al., 2017) from population genetic data summarized in the form of an allele frequency spectrum. In a typical dadi analysis, the user specifies a model with parameters that represent population sizes, migration rates, divergence times, and/or selection coefficients. For a given set of a parameters, dadi computes the expected allele frequency spectrum, from which the composite likelihood of the sample data can be calculated. Nonlinear optimization is then used to optimize parameters to maximize that likelihood, and the maximumlikelihood values are then interpreted to gain insight into past population genetic processes. During optimization, the model will be evaluated hundreds or thousands of times, leading to substantial computational expense. Here, I show that computing on Graphics Processing Units (GPUs) can massively speed dadi model computation and thus inference.

Modern graphics processing units (GPUs) provide enormous computing power for data parallel tasks, in which the same operations are applied to many entries in memory (Owens et al., 2008). But exploiting this power often demands new algorithms. For example, in computational biology there has been extensive research into GPU algorithms for sequence alignment (Manavski & Valle, 2008) and search (Vouzis & Sahinidis, 2011). In genomics, GPU algorithms have been developed for variant calling (Luo et al., 2013) and secondary analysis (Luo et al., 2014) of short-read sequencing data. But GPU computing has rarely been applied to population genetic simulation or inference. Recently, Lawrie (2017) developed a GPU implementation of the single-locus Wright-Fisher model, finding speedups of over 250 times compared to a CPU implementation. Previously, Zhou et al. (2015) implemented a subset of the IM program for inferring isolation-with-migration demographic models (Hey & Nielsen, 2004) on a GPU, demonstrating speedups of around 50 times. Here, I show that GPU computing can dramatically speedup dadi analyses.

For dadi, the limiting computation is the numerical solution of a partial differential equation (PDE) to model the dynamics of the population distribution of allele frequencies *ϕ* (Kimura, 1964). In dadi, this PDE is solved using an alternating direction implicit scheme based on the Crank-Nicolson method (Fig. 1A; Press et al. (2007)). Evolving *ϕ* forward in time then reduces to solving a large number of tridiagonal linear systems. Model parameters, including population sizes, migration rates, and selection coefficients, affect the *a, b*, and *c* diagonal vectors of these systems (Fig 1A). Unlike general matrix equations, the tridiagonal structure enables these systems to be solved in linear time, using serial Thomas algorithm (Press et al., 2007).

**Figure 1:**
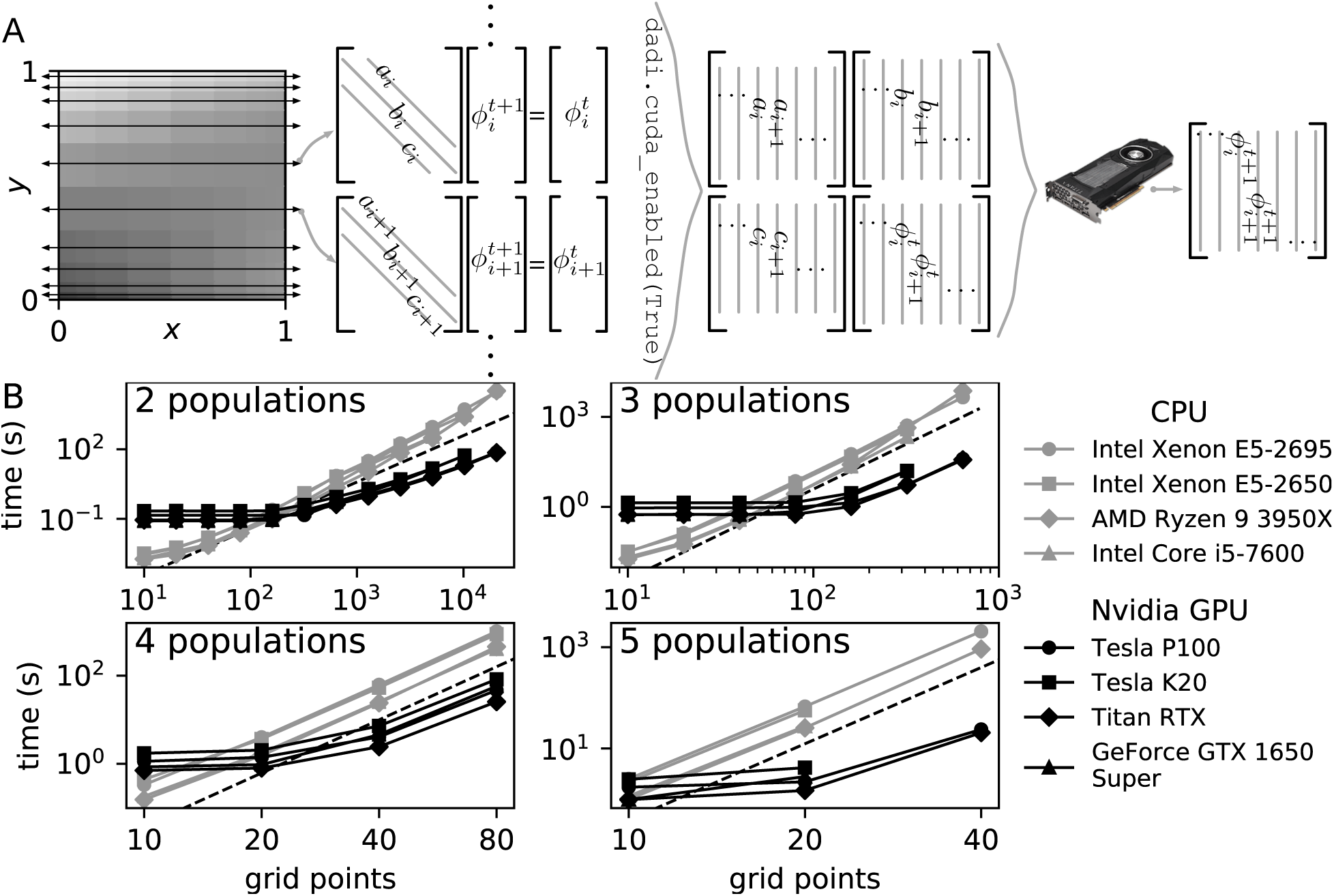
A) Illustration of the dadi numerical integration pipeline. During each timestep, the population allele density *ϕ* is propagated forward separately for each population axis. Each row in the non-uniform grid over which *ϕ* is approximated yields a tridiagonal linear system that must be solved. In the GPU implementation, the coefficients of all the systems are computed into dense matrices, and the systems are solved in parallel. B) Performance of CPU and GPU dadi implementations (smaller times are superior). For coarse simulations with few grid points, the CPU implementation is faster. But for simulations with many grid points, the GPU implementation can be orders of magnitude faster. In each panel, the dashed line indicates the approximate scaling of memory usage and total floating point operations.

Because diffusion PDEs similar to those in dadi are encountered in many fields, substantial research has been done on GPU algorithms for tridiagonal systems. Early work by Pixar employed the parallel cyclic-reduction algorithm (Hockney, 1965) to solve tridiagonal systems that arise in computer graphics on the GPU (Kass et al., 2006). Later work demonstrated that optimizing such algorithms is complex and that some parallel algorithms are numerically unstable (Zhang et al., 2010). Recently, Valero-Lara et al. (2018) showed that when the number of linear systems to be solved is large, the serial Thomas algorithm can be more efficient on the GPU than these parallel algorithms. This is because the Thomas algorithm enables more efficient memory access, if the diagonal elements *a, b*, and *c* are stored in dense matrices (Fig. 1A).

Programming GPUs demands special-purpose frameworks and libraries. Two frameworks dominate the field. The Open Computing Language (OpenCL) is a standard for writing parallel code that is portable across CPUs, GPUs, field-programmable gate arrays, and other specialized hardware (Stone et al., 2010). The Compute Unified Device Architecture (CUDA) is developed by the Nvidia Corporation specifically for use on its GPUs (Nickolls et al., 2008). In general, these frameworks offer similar performance and programming convenience (Holm et al., 2020). OpenCL is available on more platforms, particularly because CUDA is longer supported on OS X, but more libraries are available for CUDA than OpenCL. In particular, the Valero-Lara et al. (2018) algorithm for efficiently solving batches of tridiagonal systems was recently integrated into the CUDA standard library. Thus, the GPU implementation of dadi requires a CUDA-compatible GPU from Nvidia. dadi is primarily written in Python; to interface with CUDA I used the PyCUDA (Klöckner et al., 2012) and scikit-cuda (Givon et al., 2019) libraries, which can be easily installed from the Python Package Index using pip.

For the end user, dadi GPU usage is transparent, requirement only a single call to `dadi.cuda_enabled(True)`.

To evaluate performance of the dadi GPU implementation, I compared times to compute the population distribution of allele frequencies. The key parameter governing computation time is the number of grid points, *pts*, used to approximate the numerical solution of the PDE. In practice, the number of grid points must be larger than the largest data sample size, and many more points are needed when simulating strong selection (Kim et al., 2017). The *ϕ* matrix has dimensionality *P*, the number of populations modeled. So the number of elements in *ϕ* scales as *pts^P^* and so does the expected computation time. I tested models with differing numbers of populations, drawing on the stdpopsim resource where possible (Adrion et al., 2020) on multiple CPUs and GPUs. The benchmarking code is available in the dadi source repository: https://bitbucket.org/gutenkunstlab/dadi/src/master/examples/CUDA.

dadi is most often used with two- or three-population models, and in both cases the GPU implementation can be substantially faster than the CPU implementation (Fig. 1B). My two-population test model was the three-epoch model estimated for African and European Drosophila melanogaster by Li & Stephan (2006). My three-population test model was the Out-of-Africa model for humans estimated by Gutenkunst et al. (2009). The server-grade Tesla P100 GPU is over 400 times faster than the corresponding system CPU for two populations and many grid points. Even the mid-range consumer-grade GeForce GTX 1650 Super GPU is over 10 times faster than the system CPU for large 2D and 3D systems. These speed differences are illustrated more directly in Fig. S1, where the ratios of the CPU and GPU times on the same systems are shown. For all systems tested, the GPU began to outperform the CPU at around 150 grid points for two populations and at around 50 grid points for three populations, which are values regularly used with typical data analyses.

Given the dramatic speed up provided by GPU computing, I extended dadi to four- and five-population models. Tests with the four-population New World model from Gutenkunst et al. (2009) and the five-population archaic admixture model from Ragsdale & Gravel (2019) again showed that GPUs can substantially outperform CPUs.

Keys to efficient GPU computing include minimizing data transfer and managing memory usage. For dadi, the *ϕ* matrix is copied to the GPU at the outset of each integration function. All computations then remain on the GPU through potentially many time steps, until the *ϕ* matrix is copied back when the integration function exits. One subtlety is that during each time step the *ϕ* matrix must be transposed and reshaped between integration directions, to maintain the expected alignment of the *a*, *b*, *c*, and *ϕ* matrices for the batch tridiagonal solver algorithm. For example, integrating a three-population scenario with 100 grid points involves simultaneously solving 10,000 tridiagonal systems of size 100, so the naturally 100×100×100 *ϕ* array must be reshaped to 100 × 10, 000, then reshaped for the next direction of integration, and so forth. Memory is typically more limited on GPUs than on the host systems. For dadi, the full dense *a, b, c*, and *ϕ* matrices are stored as double-precision. For *P* populations and *pts* grid points, memory usage is thus roughly 4 × 8 × *pts^P^*/1024^3^ gigabytes (GB). For 3 populations, a modern mid-range GPU with 4 GB of RAM can thus scale to pts = 400, while a high-end GPU with 24 GB of RAM can scale to pts = 900. A future implementation of dadi on the GPU could potentially only compute portions of the a, b, and c matrices at a time, reducing memory usage but substantially increasing code complexity.

GPU programming can involve unusual performance decisions. For example, if parameter values are constant during an integration, then the *a, b*, and *c* matrices do not change between time steps and could be cached. Remarkably, for systems large enough to favor the GPU over the CPU, it is faster to recalculate those matrices for each time step, rather than caching them (Fig. S2). In CUDA, computational threads are organized into blocks that can share local memory, and optimizing block size can be important for maximal performance (Ryoo et al., 2008). For dadi, no communication between threads is needed, so the optimal block size is large (Fig. S3).

For many users, the ultimate benefit of GPU computing is high performance at low cost. In this respect, the performance of the mid-range consumer-grade GeForce GTX 1650 Super GPU is remarkable. As of writing, the GeForce costs roughly $200, compared to roughly $2,500 for the high-end consumer-grade Titan RTX and roughly $6,000 for the server-grade Tesla P100. Yet the performance of the GeForce is within a factor of three of the high-end GPUs (Fig. S4), even though the P100 theoretically has a 30-fold advantage in double-precision operations per second. This suggests that the dadi workload is bound by memory bandwidth rather than arithmetic. It also shows that the massive benefits of GPU computing for dadi are easily accessible to end users.

GPU computing offers substantial performance benefits for dadi, with minimal user burden. These performance improvements increase dadi’s competitiveness with alternative methods for calculating the allele frequency spectrum, such as moments (Jouganous et al., 2017). They also make analysis of more complex models, including those with four and five populations, computationally feasible. Lastly, the large benefits of even consumer-grade GPUs for dadi suggest that GPU computing may also be worth considering for other software in computational population genetics.

## Acknowledgments

This work was supported by the National Institute of General Medical Sciences of the National Institutes of Health (R01GM127348 to R.N.G.). This material is based upon High Performance Computing (HPC) resources supported by the University of Arizona TRIF, UITS, and Research, Innovation, and Impact (RII) and maintained by the UArizona Research Technologies department. I thank Xin Huang for benchmarking assistance and Andreas Klöckner for guidance to the Python CUDA ecosystem.

---

**Figure S1:**
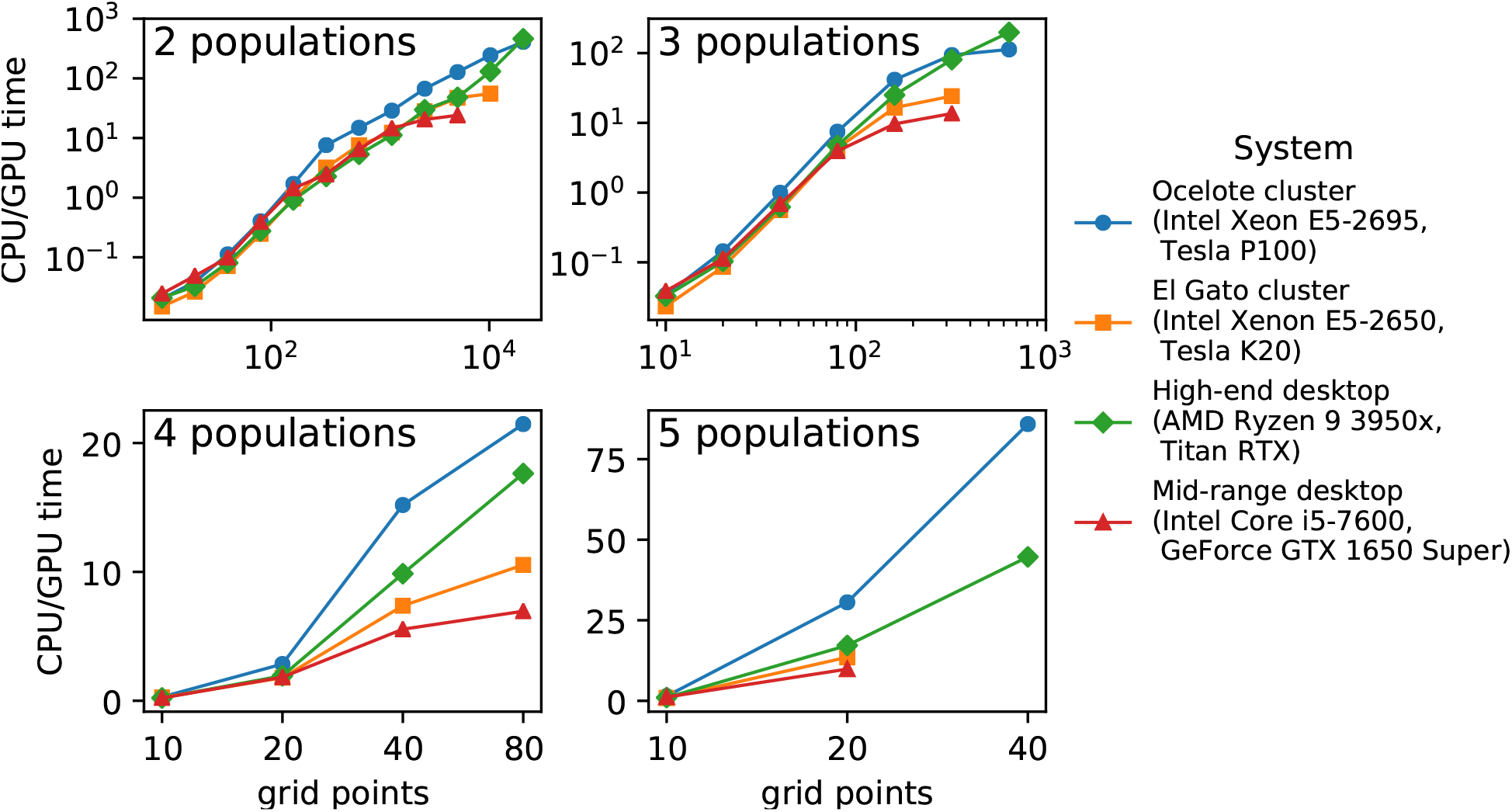
GPU vs CPU speed ups. The results from Fig. 1B are replotted, now showing the ratio between CPU and GPU times on the same systems. The Ocelote and El Gato clusters are University of Arizona centralized Higher Performance Computing resources.

**Figure S2:**
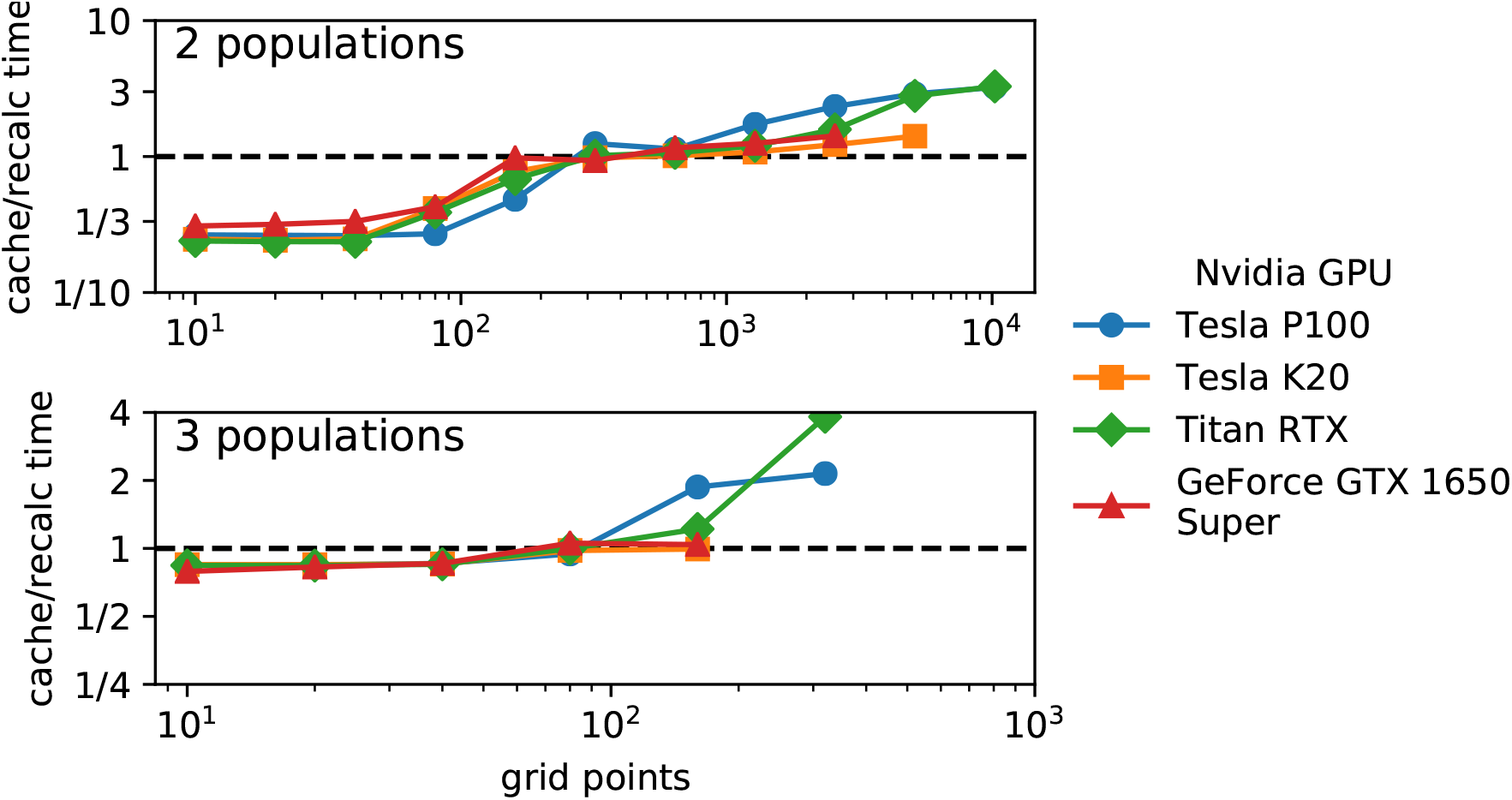
Caching vs recalculating *a, b*, and *c* matrices. For the two-population test model and a modified three-population model that uses piecewise constant parameters, plotted is the ratio of times for implementations that cache a, b, and c matrices or recalculate them each time step. Recalculating each time step is faster for larger grid point settings.

**Figure S3:**
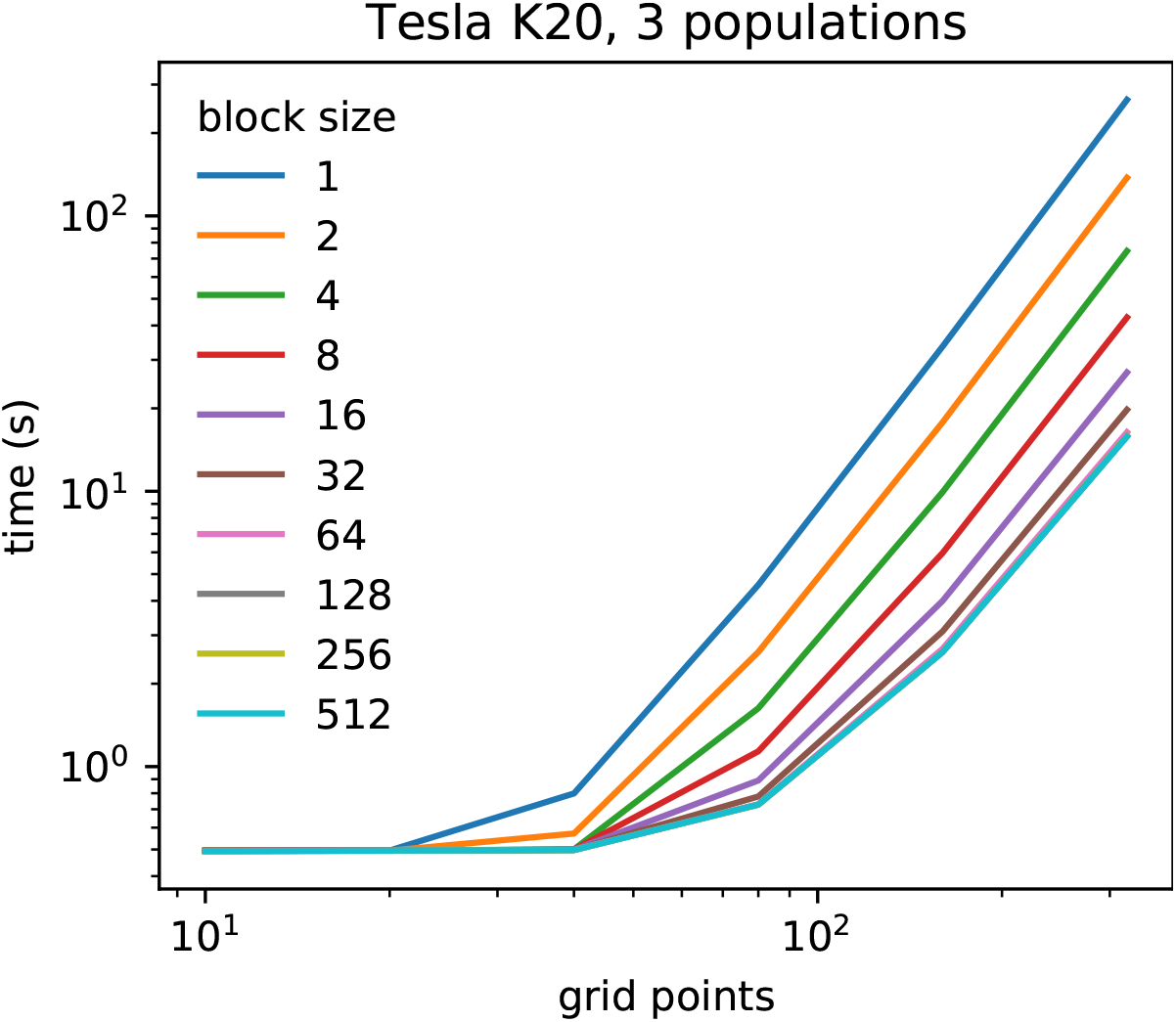
Performance versus thread block size. For the three-population test model and a Tesla K20 GPU, plotted is the integration time versus grid points for multiple block size settings. Large blocksizes are optimal.

**Figure S4:**
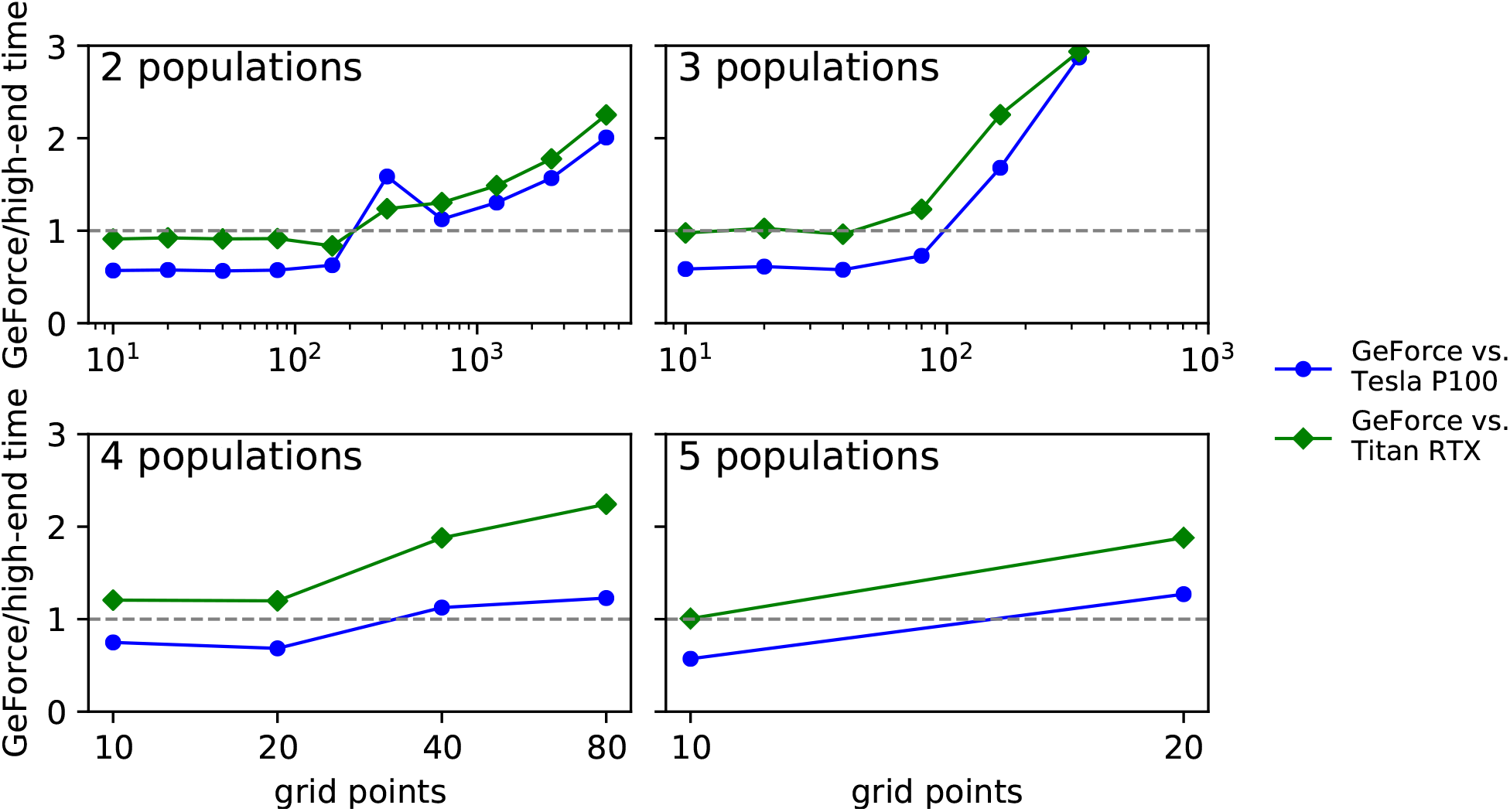
Performance of inexpensive vs expensive GPUs. For the test models, plotted is the ratio of the computation time of the GeForce GTX 1650 Super GPU to the time of two much more expensive GPUs. The advantage of the more expensive GPUs grows with problem size, but in these tests it never exceeds a factor of three.

## References

Adrion JR, Cole CB, Dukler N, Galloway JG, Gladstein AL, Gower G, Kyriazis CC, Ragsdale AP, Tsambos G, Baumdicker F, Carlson J, Cartwright RA, Durvasula A, Gronau I, Kim BY, McKenzie P, Messer PW, Noskova E, Ortega Del Vecchyo D, Racimo F, Struck TJ, Gravel S, Gutenkunst RN, Lohmueller KE, Ralph PL, Schrider DR, Siepel A, Kelleher J, Kern AD (2020) A community-maintained standard library of population genetic models. eLife 9:e54967.

Givon LE, Unterthiner T, Erichson NB, Chiang DW, Larson E, Pfister L, Dieleman S, Lee GR, van der Walt S, Menn B, Moldovan TM, Bastien F, Shi X, Schluter J, Thomas B, Capdevila C, Rubinsteyn A, Forbes MM, Frelinger J, Klein T, Merry B, Merill N, Pastewka L, Liu LY, Clarkson S, Rader M, Taylor S, Bergeron A, Ukani NH, Wang F, Lee WK, Zhou Y (2019) scikit-cuda 0.5.3: a Python interface to GPU-powered libraries. http://dx.doi.org/10.5281/zenodo.3229433.

Gutenkunst RN, Hernandez RD, Williamson SH, Bustamante CD (2009) Inferring the joint demographic history of multiple populations from multidimensional SNP frequency data. PLoS Genetics 5:e1000695.

Hey J, Nielsen R (2004) Multilocus Methods for Estimating Population Sizes, Migration Rates and Divergence Time, With Applications to the Divergence of Drosophila pseudoobscura and D. persimilis. Genetics 167:747.

Hockney RW (1965) A Fast Direct Solution of Poisson’s Equation Using Fourier Analysis. Journal of the ACM (JACM) 12:95.

Holm HH, Brodtkorb AR, Sætra ML (2020) GPU computing with python: Performance, energy efficiency and usability. Computation 8:4.

Jouganous J, Long W, Ragsdale AP, Gravel S (2017) Inferring the Joint Demographic History of Multiple Populations: Beyond the Diffusion Approximation. Genetics 206:1549.

Kass M, Lefohn A, Owens J (2006) Interactive Depth of Field Using Simulated Diffusion on a GPU. Technical report, Pixar.

Kim BY, Huber CD, Lohmueller KE (2017) Inference of the distribution of selection coefficients for new nonsynonymous mutations using large samples. Genetics 206:345.

Kimura M (1964) Diffusion Models in Population Genetics. Journal of Applied Probability 1:177.

Klöockner A, Pinto N, Lee Y, Catanzaro B, Ivanov P, Fasih A (2012) PyCUDA and PyOpenCL: A scripting-based approach to GPU run-time code generation. Parallel Computing 38:157.

Lawrie DS (2017) Accelerating Wright-Fisher forward simulations on the graphics processing unit. G3: Genes, Genomes, Genetics 7:3229.

Li H, Stephan W (2006) Inferring the demographic history and rate of adaptive substitution in Drosophila. PLoS Genetics 2:1580.

Luo R, Wong T, Zhu J, Liu CM, Zhu X, Wu E, Lee LK, Lin H, Zhu W, Cheung DW, Ting HF, Yiu SM, Peng S, Yu C, Li Y, Li R, Lam TW (2013) SOAP3-dp: Fast, Accurate and Sensitive GPU-Based Short Read Aligner. PLoS ONE 8:e65632.

Luo R, Wong YL, Law WC, Lee LK, Cheung J, Liu CM, Lam TW (2014) BALSA: Integrated secondary analysis for whole-genome and whole-exome sequencing, accelerated by GPU. PeerJ 2:e421.

Manavski SA, Valle G (2008) CUDA compatible GPU cards as efficient hardware accelerators for Smith-Waterman sequence alignment. BMC Bioinformatics 9:S10.

Nickolls J, Buck IaN, Garland M (2008) Scalable Parallel with CUDA. Queue 6:40.

Owens JD, Houston M, Luebke D, Green S, Stone JE, Phillips JC (2008) GPU computing. Proceedings of the IEEE 96:879.

Press WH, Teukolsky SA, Vetterling WT, Flannery BP (2007) Numerical Recipes 3rd Edition: The Art of Scientific Computing. Cambridge University Press, 3 edition.

Ragsdale AP, Gravel S (2019) Models of archaic admixture and recent history from two-locus statistics. PLoS Genetics 15:e1008204.

Ryoo S, Rodrigues CI, Baghsorkhi SS, Kirk DB, Hwu WmW, Stone SS (2008) Optimization Principles and Application Performance Evaluation of a Multithreaded GPU Using CUDA. In PPoPP ‘08 Proceedings of the 13th ACM SIGPLAN Symposium on Principles and practice of parallel programming, pages 73–82.

Stone JE, Gohara D, Shi G (2010) A Parallel Programming Standard for Heterogeneous Computing Systems. Computing in Science & Engineering 12:66.

Valero-Lara P, Martinez-Pérez I, Sirvent R, Martorell X, Pena AJ (2018) cuThomasBatch and cuThomasVBatch, CUDA Routines to compute batch of tridiagonal systems on NVIDIA GPUs. Concurrency Computation 30:e4909.

Vouzis PD, Sahinidis NV (2011) GPU-BLAST: Using graphics processors to accelerate protein sequence alignment. Bioinformatics 27:182.

Zhang Y, Cohen J, Owens JD (2010) Fast tridiagonal solvers on the GPU. ACM SIGPLAN Notices 45:127.

Zhou C, Lang X, Wang Y, Zhu C (2015) gPGA: GPU accelerated population genetics analyses. PLoS ONE 10:e0135028.

